# SARS-CoV-2 infection disrupts syncytial and endothelial integrity and alters PLGF levels in the placenta

**DOI:** 10.1101/2025.06.27.661568

**Authors:** Brittany R. Jones, Guilherme M. Nobrega, Deepak Kumar, Emily Diveley, Arthur Antolini-Tavares, Renato T Souza, José Guilherme Cecatti, Jeannie C. Kelly, Maria Laura Costa, Indira U. Mysorekar

## Abstract

**Introduction:** SARS-CoV-2 infection during pregnancy has been associated with an increased risk for several pregnancy-related disorders, particularly preeclampsia (PE). However, there are limited studies determining the impact of SARS-CoV-2 on placental physiology and function.

**Methods:** Placental samples were acquired from two large prospective cohorts: STOP-COVID19 and REBRACO studies. Placental villous tissues (VTs) were collected from pregnant women who tested positive for SARS-CoV-2 without PE during pregnancy. Immunohistochemistry and immunofluorescence were used to assess pathological features known to be altered in PE, including 1) syncytial knot formation; 2) alterations in renin-angiotensin system components; 3) and endothelial integrity. Maternal serum was collected to examine AT_1_ autoantibodies levels using an immunoassay.

**Results:** SARS-CoV-2 viral proteins spike, nucleocapsid, and ORF3a were observed in the syncytiotrophoblast layer and stroma of placental VT. SARS-CoV-2-infected placentas exhibited increased numbers of syncytial knots, which were positive for Flt-1 and SARS-CoV-2 viral proteins. In addition, the presence of placental infarctions and excessive fibrin deposits was also observed in infected placentas. Infection was associated with decreased placental expression of PlGF and an increase in the placental Flt-1/PlGF expression ratio, mostly driven by PlGF. No significant changes in maternal serum AT_1_AA levels were observed. Finally, SARS-CoV-2-infected placentas exhibited a significant decrease in vimentin expression.

**Discussion:** SARS-CoV-2 infection negatively impacts placental integrity in the form of increased syncytial knots, dysregulated RAS components, and endothelial damage. Since all these features are similarly disrupted in PE, this could be a mechanism through which SARS-CoV-2 infection during pregnancy increases the risk of a PE-like syndrome.

## 1. Introduction

In 2019, the outbreak of severe acute respiratory syndrome coronavirus 2 (SARS-CoV-2) infection sparked a global pandemic (COVID-19) that has resulted in the deaths of more than 7 million people worldwide [1]. There is still limited understanding of disease development and its immediate and lasting effects, especially concerning how SARS-CoV-2 infection during pregnancy impacts both pregnant women and their fetuses[2]. Pregnant women are at risk of developing severe disease outcomes, higher risk composite morbidity, and pregnancy complications, including preterm birth, stillbirth, and preeclampsia (PE) [3–8]. Although transmission from mother to fetus is rare, infection during pregnancy has been associated with placental pathology and fetal/neonatal neurodevelopmental changes [1,9–11]. Histopathological changes in the placenta as a result of SARS-CoV-2 infection have been associated with adverse pregnancy outcomes [5,12].

A significant increase in hypertensive disorders in pregnancies complicated by SARS-CoV-2 infection has been noted [13–15] with some studies suggesting a link between the virus and increased PE risk in infected pregnancies [13,16,17]. Pregnant women with moderate to severe COVID-19 infection are eight times more likely to develop early-onset PE [18,19]. This has led to the identification of a PE-like syndrome in infected pregnancies, characterized by similarities in the pathophysiology and clinical presentation of PE and SARS-CoV-2 infection during pregnancy [13,15,20–23].

PE is a pathophysiological condition where vascular resistance is abnormally increased and contributes to maternal hypertension. PE affects approximately 2-8% of pregnancies worldwide and is one of the main causes of maternal and perinatal morbidity and mortality [13,20]. Pathologically, PE is associated with aberrant increases in syncytial knots (aggregates of fused cells at the surface of the terminal villi), an indicator of maternal vascular malperfusion and maternal hypertensive disorders [24–26]. Syncytial knots reflect villous maturity and in normal pregnancies are found in ∼30% of terminal villi [27,28]; however, they are almost ubiquitous in the placentas of preeclamptic women and associated with pathologies such as decidual arteriopathy and accelerated villous maturation [27,29,30]. PE is also associated with endothelial damage, vasoconstriction, and vascular inflammation [20,31,32]. A key process dysregulated in PE is the placental renin-angiotensin system (RAS), which plays an important role in normal pregnancy by regulating placental angiogenesis, protection against infection, and promoting fetal development [33,34].

Multiple studies, including from our group, have shown SARS-CoV-2 infects multiple compartments of the placenta, particularly the villous syncytiotrophoblast cell layer, fetal macrophages, chorionic and amniotic membranes at different stages of pregnancy [17,35]. Severe SARS-CoV-2 infection has been associated with the occurrence of endothelial cell injury and platelet stress, thereby disrupting the homeostatic balance in the vasculature and thrombosis in multiple organs such as lung, kidneys, and heart [36,37].

The receptor for SARS-CoV-2 in host cells is angiotensin-converting enzyme 2 (ACE2), a key component in the activation of the proinflammatory and anti-inflammatory pathway of RAS [38–40]. We and others have shown that ACE2 is expressed in placental cells. SARS-CoV-2 infection is associated with decreased ACE2 in trophoblast cells and imbalance of RAS resulting in hyperactivation of the pro-inflammatory pathway (AT1) that leads to increased production of soluble fms-like tyrosine kinase-1 (sFlt-1/Flt-1), an anti-angiogenic protein which causes vasoconstriction and endothelial damage that may affect fetal growth and a decrease in pro-angiogenic placental growth factor (PlGF) [39,41,42]. PlGF is an important molecule during pregnancy that affects feto-placental circulation and supports trophoblast growth [43]. The balance between sFlt-1/PlGF is important for normal pregnancy, as a high plasma sFlt1/PlGF ratio is a strong predictor for risk of developing PE during pregnancy[44]. Previously, we have demonstrated that SARS-CoV-2 spike protein binding to ACE2 leads to elevated levels of sFlt-1, a signatory marker of PE [17]. Sera from SARS-CoV-2 exposed pregnant women also showed elevated levels of autoantibodies to angiotensin II type 1-receptor (AT1R-AA) prior to delivery, another signatory marker of PE and dysregulated RAS activity [17]. Histopathological analysis has shown that pregnant women infected with SARS-CoV-2 present similar placental pathology features seen in a preeclamptic placenta [13,21,36,45]. Infection can result in chorangiosis, high-grade villitis with trophoblastic destruction, fibrin deposition, and acute chorioamnionitis in the placenta of pregnant women [36].

These findings suggest a cellular and molecular pathological link between SARS-CoV-2 infection and PE. However, a comprehensive histopathological analysis of a PE-like syndrome in the context of SARS-CoV-2 infection has yet to be conducted. In this study, we aimed to examine the impact of SARS-CoV-2 on placental pathology, syncytial knot formation, and renin-angiotensin system dynamics. We compared placental samples from women exposed to SARS-CoV-2 during pregnancy, without PE, across two separate cohorts, to explore the mechanisms underlying these pathological changes. Our results indicate that SARS-CoV-2 infection disrupts placental integrity, characterized by increased syncytial knots, dysregulated RAS components, and endothelial damage.

## 2. Methods

### 2.1. Participants and biological samples

This study is a case series of placental samples collected from two large prospective cohorts: Safety, Testing/Transmission, and Outcomes in Pregnancy with COVID-19 (STOP-COVID19) collected in St. Louis, USA (December 2020 to July 2022) and the Brazilian Network of COVID-19 in Obstetrics (REBRACO) multicentric study group from participants enrolled in the coordinating center in Campinas, São Paulo, Brazil (May 2020 to June 2021) [17,32,46]. Inclusion criteria were term births, singletons, unvaccinated against SARS-CoV-2, and with placental villous tissue (VT) samples collected for analysis. Exclusion criteria included pre-term birth, pre-eclampsia, eclampsia, and stillbirths. Placental tissues were collected from pregnant women who tested positive for SARS-CoV-2 (mild to severe cases) or negative. For both cohorts, VT was collected at delivery. Maternal peripheral blood serum was collected at delivery for the REBRACO cohort.

### 2.2. Clinical Characteristics

Medical charts and study interviews were reviewed to retrieve information on general characteristics, comorbidities, maternal outcomes, and pregnancy and perinatal outcomes. General characteristics included age (≤24, 24-35, ≥36), race/ethnicity [White, Black/African American, Asian, Latinos, Native American, Multiracial (Pardo)]. In the STOP-COVID19 study, race and ethnicity were classified following the criteria of the National Institute of Health (NIH). The NIH describes race and ethnicity as Black or African American, Hispanic or Latino, Asian, American Indian or Alaska Native, Native Hawaiian or Other Pacific Islander, Multiracial, and White (Supplementary Table 2). The REBRACO considered the IBGE criteria for racial and ethnicity status. The IBGE classifies the Brazilian population into five categories based on skin color by asking participants to self-identify as White, Black, “Pardo” (brown), Yellow (East Asian), or Indigenous[47]. Comorbidities including diabetes mellitus, chronic hypertension, obesity, asthma, anemia, STORCH infections, or others (thyroid disease, alopecia areata, Sjogren’s syndrome, chronic kidney disease, cardiovascular disease, sickle cell anemia, and epilepsy) were reported. For maternal outcomes, the variables included were confirmed SARS-CoV-2 diagnosis (yes/ no, and gestational trimester at diagnosis), symptomatic SARS-CoV-2 infection, and SARS-CoV-2 epidemiological strain (Wuhan/WT, Delta, Gamma, and Omicron). SARS-CoV-2 infection was confirmed by RT-qPCR using upper respiratory secretion samples (nasopharyngeal swabs) or saliva. For pregnancy and perinatal outcomes, delivery route (C-section, vaginal) and Apgar 5 minutes score < 7 were considered.

### 2.3. H&E Staining and Immunohistochemistry (IHC)

Formalin-fixed paraffin embedded placental VT samples were sectioned at 5 μm thickness and were sequentially rehydrated using ethanol-water gradients. Sections were stained with Harris hematoxylin for 3 minutes and briefly rinsed in deionized water before destaining in 1% acid alcohol and soaked in 1% sodium bicarbonate for 5 minutes. Sections were counterstained with eosin, dehydrated using ethanol water-gradients and mounted with Permount mounting media. Slides were visualized using a 3D HISTECH slide viewer (3DHISTECH, Ltd).

### 2.4. Quantification of Syncytial Knots

Syncytial knots were quantified in H&E-stained sections using a method based on a previous study [30]. The primary investigator (BJ) was blinded to COVID-19 diagnosis and clinical characteristics of participants. Syncytial knots were defined as a multilayered aggregate of at least 5 syncytiotrophoblastic nuclei protruding from the villous surface that were not in direct contact with any adjacent villi. Syncytial knots were only included if they were, I) suspended between the intervillous space, or II) located on the edge of the syncytium. The number of syncytial knots within four individual 4 mm^2^ quadrants per slide were counted and averaged. Any syncytial aggregates connecting two villi were excluded.

### 2.5. Immunofluorescence analysis

The primary investigator (BJ) was blinded to COVID-19 diagnosis and clinical characteristics of participants during IF staining. Antigen retrieval was performed on VT sections as described for IHC, except for staining for SARS-CoV-2 viral proteins. Sections were washed with PBS and blocked with 5% horse serum with 0.3% Triton X for 1 hour at room temperature. After blocking, tissue sections were incubated overnight at 4 °C with primary antibodies specific for SARS-CoV-2 Nucleocapsid (1:300, Invitrogen), Spike (1:500, Invitrogen), ORF3a (1:200, R&D Systems), Cytokeratin 5 (1:500, Abcam) FLT1 (1:200, R&D Systems), PlGF (1:100, biorbyt), and Vimentin (1:100, NOVUS). Following washing (PBS with 0.1% Triton X), sections were stained with Alexa Fluor-conjugated secondary antibodies at room temperature for one hour: anti-chicken (1:200, Invitrogen), anti-mouse (1:500, Invitrogen), anti-rabbit (1:200 and 1:500, Invitrogen), and anti-goat (1:300, ThermoFisher). Following additional washing, autofluorescence was quenched using the Vector TrueView Autofluorescence Quenching Kit (Vector Laboratories, CA). Sections were counterstained with Hoescht dye before imaging with a Nikon AX-R Point Scanning Confocal microscope (Nikon, NY) at 20X, 40X, and 60X. For quantification, the mean fluorescence intensity of four random fields at 20X was averaged for each slide.

### 2.6. Angiotensin II Type 1-Receptor Autoantibody (AT_1_-AA) ELISA

Levels of AT_1_AA were measured in maternal peripheral serum from uninfected and SARS-CoV-2-infected pregnant women using a commercial ELISA kit (MyBiosource). All samples were run in duplicate as per manufacturer’s instruction and read using a microplate reader at 450 nm. The author was blinded to COVID-19 diagnosis and clinical characteristics of participants.

### 2.7. Data Analysis

Cases of both cohorts were merged and classified into two groups: no diagnosis of SARS-CoV-2 infection during pregnancy, SARS-CoV-2(-); and positive SARS-CoV-2 diagnosis at any point during pregnancy, SARS-CoV-2(+). The number of samples for SARS-CoV-2(-) and SARS-CoV-2(+) groups were determined based on the minimum detectable effect, given a Type 1 error rate of 0.05 and a desired statistical power of 80%. Normality tests were performed on sample data prior to analysis. Comparisons between groups for categorical variables were performed using Odds Ratio with 95% confidence interval (CI), and uncorrected Chi-square test or Fisher’s exact (expected results less than 5) for clinical characteristics using EPInfo (CDC, v7.2.5 2021). Statistical analysis for continuous variables was performed using nonparametric Mann-Whitney U test for IHC, IF, and ELISA experiments using Graphpad Prism 10.3.0. P < 0.05 was considered statistically significant.

## 3. Results

### 3.1. Clinical Characteristics

Details of the demographics for the STOP-COVID19 (n = 38) and REBRACO (n = 38) cohorts are shown in Supplementary Table 1. We note no significant differences in clinical characteristics including pregnancy and perinatal outcomes, most of the comorbidities, and age (p-value=0.2547) between the STOP-COVID19 and REBRACO cohorts. We saw significant differences in maternal outcomes as SARS-CoV-2 infection presented more in the 1^st^ trimester in the STOP-COVID19 (p-value<0.0001) cohort and primarily in the 3^rd^ trimester in the REBRACO cohort (Supplementary Table 1). There was no difference in 2^nd^ trimester infections between the two cohorts, however, the STOP-COVID19 study consisted primarily of SARS-CoV-2 strains Delta and Omicron (p-value<0.0001) whereas the REBRACO study participants were primarily infected with Wuhan and Gamma strains (p-value<0.0001). Race could not be compared because the USA and Brazil classify race/ethnicity differently. Given the lack of difference in most comorbidities, age, and pregnancy outcomes, we combined both cohorts when evaluating demographic differences between SARS-CoV-2(-) and SARS-CoV-2(+) groups (Table 1). When the two cohorts were combined, maternal age, comorbidities, and pregnancy outcomes were not significantly different between SARS-CoV-2(-) and SARS-CoV-2(+) cases. The exceptions to this were obesity, which was higher in the SARS-CoV-2(+) group (p-value =0.0075), and anemia (p-value =0.0123), which was higher in the SARS-CoV-2(-) group (Table 1).

**Table 1:** Sociodemographic data, comorbidities, maternal, and neonatal outcomes for SARS-CoV-2 negative (n=29) and SARS-CoV-2 positive (n=47) cases.

| Variable | SARS-CoV-2 (-) |  | SARS-CoV-2 (+) |  | Odds Ratio (95% CI) | *p-value* |
| --- | --- | --- | --- | --- | --- | --- |
| *N* | 29 |  | 47 |  |  |  |
| **General Characteristics** |  |  |  |  |  |  |
| Age |  |  |  |  |  |  |
| ≤ 24 | 4 | 13.8% | 12 | 25.5% | - | 0.3325** |
| 24 - 35 | 21 | 72.4% | 26 | 55.3% | - |  |
| ≥ 36 | 4 | 13.8% | 9 | 19.1% | - |  |
| **Comorbidities** |  |  |  |  |  |  |
|  | (n = 27) |  | (n = 42) |  |  |  |
| Diabetes Mellitus | 4 | 14.8% | 10 | 23.8% | 1.7969 (0.5009 ~ 6.4456) | 0.5412** |
| Chronic Arterial Hypertension | 4 | 14.8% | 9 | 21.4% | 1.5583 (0.4311 ~ 6.4495) | 0.5469** |
| Obesity | 6 | 22.2% | 23 | 54.8% | 4.2368 (1.4170 ~ 12.6265) | **0.0075\*** |
| Asthma | 5 | 18.5% | 2 | 4.8% | 0.2200 (0.0394 ~ 1.2291) | 0.1018** |
| Anaemia | 6 | 22.2% | 1 | 2.4% | 0.0854 (0.0096 ~ 0.7562) | **0.0123\*** |
| STORCH | 1 | 3.7% | 5 | 11.9% | 3.5135 (0.3874 ~ 31.8647) | 0.3922** |
| Others | 2 | 7.4% | 4 | 9.5% | 1.3158 (0.2239 ~ 7.7315) | 1.0000** |
| **Maternal Outcomes** |  |  |  |  |  |  |
|  | (n = 29) |  | (n = 47) |  |  |  |
| Hemorrhage | 0 | - | 3 | 5.7% | - | 0.2826** |
| **Pregnancy and Perinatal Outcomes** |  |  |  |  |  |  |
| Delivery Route |  |  |  |  |  |  |
| C-section | 8 | 24.2% | 21 | 44.7% | 2.1202 (0.7824 ~ 5.7455) | 0.1362* |
| Vaginal | 21 | 63.6% | 26 | 55.3% |  |  |
| Apgar 5 minutes < 7 | 1 | 3.0% | 0 | - | - | 0.3816** |

Others: thyroid disease, alopecia areata, Sjogren's syndrome, chronic kidney disease, cardiovascular disease, sickle cell anemia, and epilepsy. \*Chi-Square \*\* Fisher's

### 3.2. Placentas from SARS-CoV-2 infection during pregnancy are positive for viral proteins

Within the infected cohort, we observed localized foci of SARS-CoV-2 viral proteins spike and nucleocapsid in the STB layer and stroma in the chorionic villi (Figure 1A, Inset A and B), consistent with studies that have shown SARS-CoV-2 infects multiple compartments of the placenta[17,35,48]. SARS-CoV-2 accessory protein, ORF3a was observed in the STB layer of placental sections from infected pregnancies, indicative of active viral replication (Figure 1B, Inset C and D). Additionally, we observed colocalization of spike and ORF3a in the STB layer (Figure 1C, Inset E and F) in placental VT.

**Figure 1:**
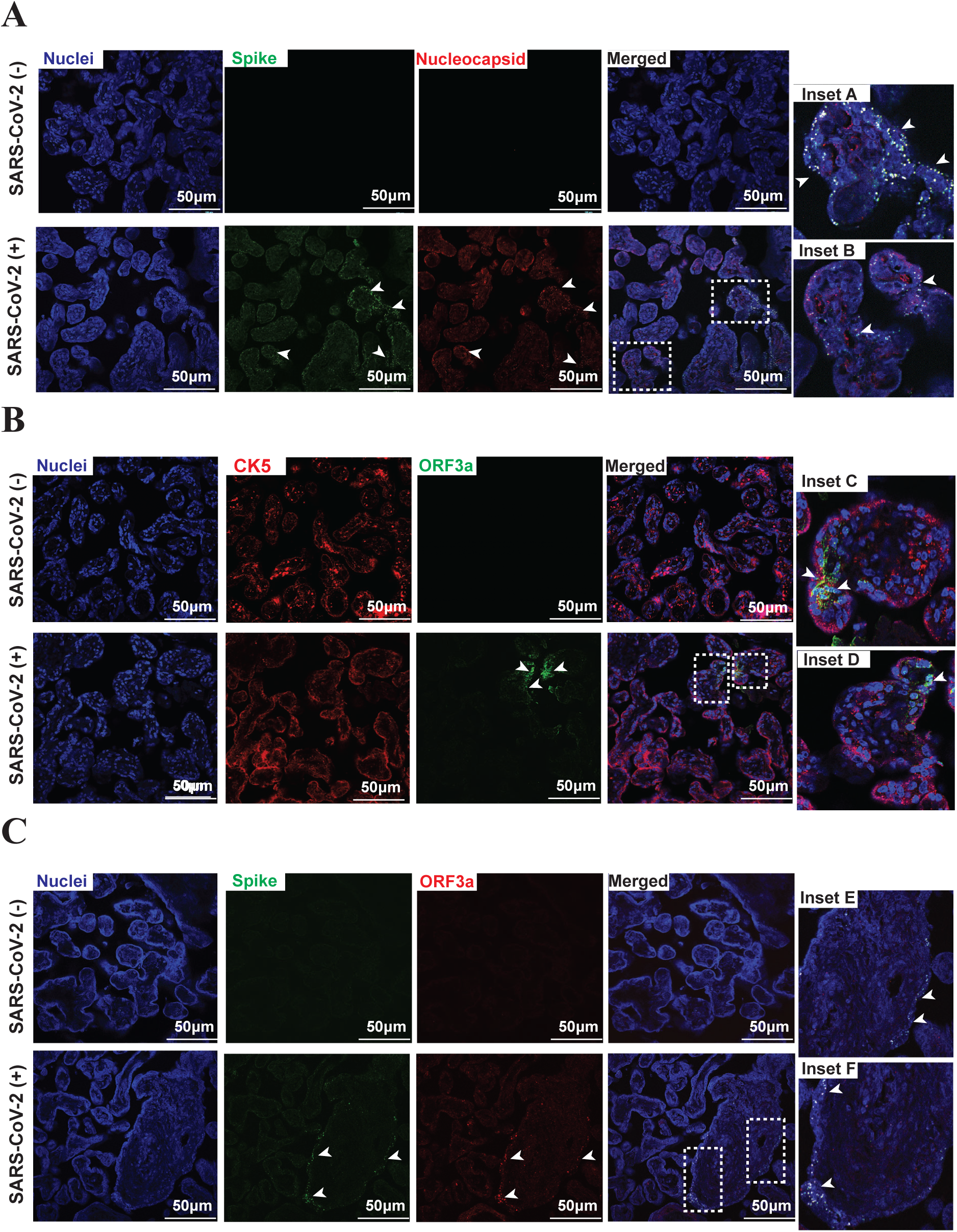
Foci of SARS-CoV-2 infection in the placenta. (A) Detection of SARS-CoV-2 proteins spike (green), nucleocapsid (red), and colocalization (inset A and B) in villous tissue from SARS-CoV-2 positive term placentas (magnification scale 20X; scale bar represents 50 µm), Nuclei (blue). (B) Presence of SARS-CoV-2 ORF3a (green) in STB layer (Inset C and D); STB layer outlined with CK5 (red) (magnification scale 20X; scale bar represents 50 µm). (C) SARS-CoV-2 proteins spike (green) and ORF3a (red) detected in villous tissue and colocalization of (Inset E and F) of viral proteins in STB layer (magnification scale 20X; scale bar represents 50 µm).

### 3.3. SARS-CoV-2 infection is associated with increased syncytial knot formation

Syncytial knots were observed in both uninfected and SARS-CoV-2-infected placental sections (Figure 2A) localized on the edge of the villi (Inset A, C, and D) or suspended between villi (Inset B, E, and F). However, the number of syncytial knots was significantly increased in SARS-CoV-2-infected VT (p-value<0.0001, Figure 2B). Syncytial knots were found to express SARS-CoV-2 spike, nucleocapsid, and ORF3a proteins (Figure 1, Inset A and Inset C). Placental sections were also examined for other pathological markers or damage using histology. In addition to increased syncytial knots, placentas from infected pregnancies exhibited placental infarctions and excessive fibrin deposition, suggesting placental damage (Figure 1C and 1D).

**Figure 2:**
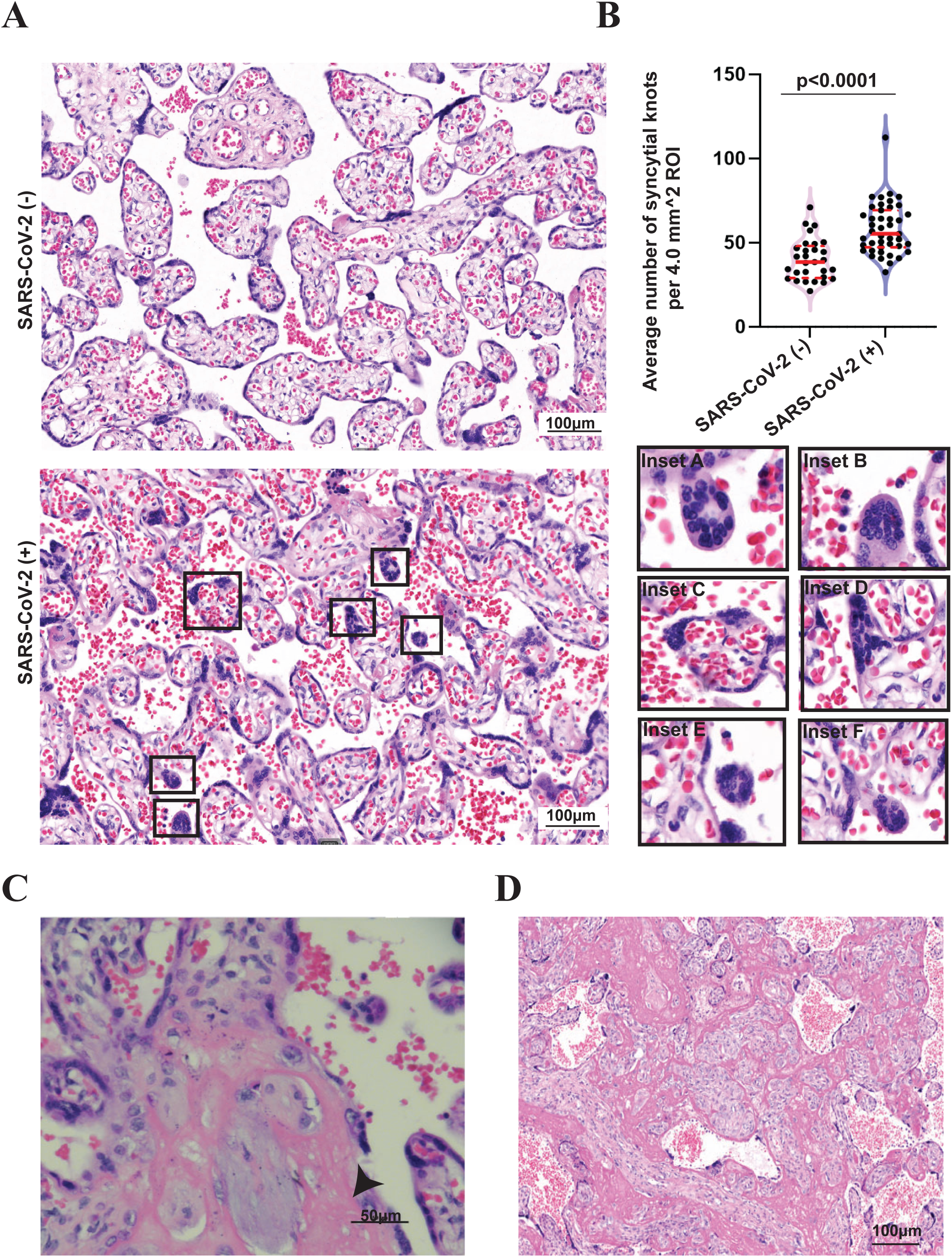
Increased formation of syncytial knots in SARS-CoV-2 infected placentas. (A) Presence of SK in SARS-CoV-2 positive placentas. Syncytial knots were located in the maternal circulation (Inset A, C, and D) or suspended between villi (Inset B, E, and F). (B) Semiquantitive analysis of syncytial knots formation in uninfected and SARS-CoV-2 infected placentas. Values are expressed as mean +/-standard error of the mean (SEM); * p <0.05, ** p < 0.01, ***: p < 0.001; ns: non-significant. (C-D) Presence of placental infarction (C), and excessive fibrin deposition (D) in SARS-CoV-2 infected placentas.

### 3.4. SARS-CoV-2 infection is associated with decreased PlGF

We previously reported a significant decrease in ACE2 expression in SARS-CoV-2-infected placentas, suggesting that placental RAS is dysregulated by infection[17]. In our current study, we observed the presence of Flt-1 and PlGF in the STB layer with puncta staining in the stroma in both the uninfected and infected placental sections (Figure 3). Flt-1 was also detected in syncytial knots (Figure 3A, Inset A and B), including in knots suspended between villi (Figure 3A Inset B). Further, SARS-CoV-2-infected placentas showed a significant decrease in expression of PlGF, mainly in focal areas of the VT (Figure 3A and 3B; p-value<0.0001). While there was no significant difference in Flt-1 expression between the two groups (Figure 3B, p-value=0.0664), the placental Flt-1/PlGF ratio was significantly higher in SARS-CoV2 (+) placentas (Figure 3C; p-value=0.0002), suggesting dysregulation of RAS. Next, we evaluated the levels of AT_1_AA in the maternal serum (Figure 3D). We saw no change in the levels of AT_1_AA (p-value=0.6484). Together, our findings demonstrate SARS-CoV-2 infection leads to reduced expression of PlGF, resulting in a higher placental Flt-1/PlGF ratio.

**Figure 3:**
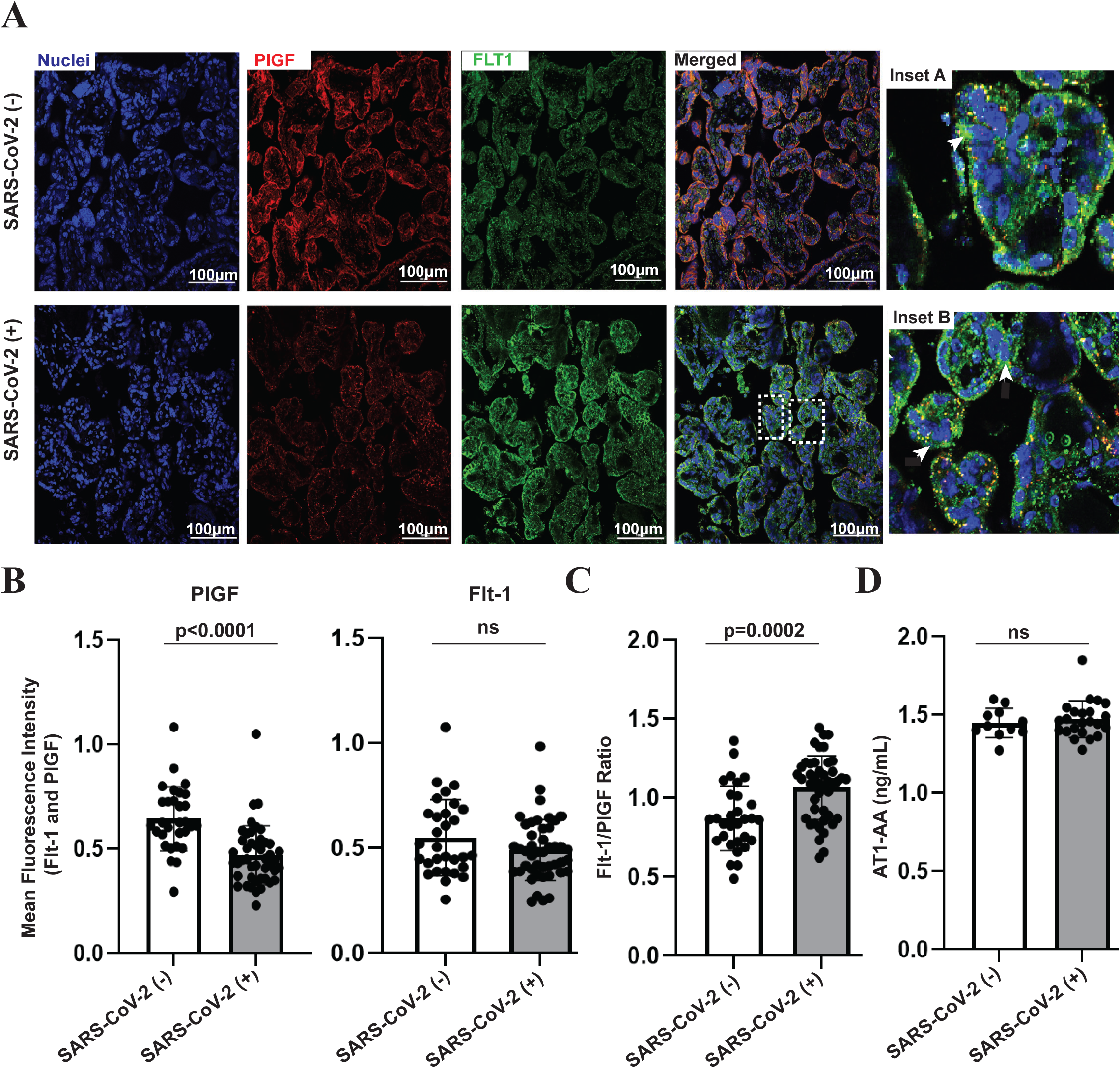
SARS-CoV-2 infection is associated with changes in RAS components in the placenta. (A) Immunofluorescence images of uninfected and SARS-CoV-2 infected villous tissue showing expression of PlGF (red), Flt-1 (green) (magnification scale 20X; scale bar represents 100 µm). Presence of Flt-1 in SK of SARS-CoV-2-infected placentas (Inset A and B). (B-C) Quantification of mean fluorescence intensity of PlGF and Flt-1 in uninfected and infected placentas (B) and Flt-1/PlGF ratio (C). (D) Pre-delivery of angiotensin II type 1-receptor autoantibody (AT_1_-AA) levels in sera of uninfected and SARS-CoV-2-infected pregnant women. Bars represent means and error bars represent standard error of the mean (SEM); *: * p <0.05, ** p < 0.01, ***: p < 0.001; ns: nonsignificant difference.

### 3.5. SARS-CoV-2 infection is associated with endothelial injury in the placenta

Severe SARS-CoV-2 infection can lead to endothelial injury, causing excessive platelet stress, inflammation, and disruption of the homeostatic balance in the vasculature, as seen in the lungs of COVID-19 patients [37]. We examined SARS-CoV-2-infected placental VT sections and noted morphological changes, including chorangiosis, suggesting vascular changes in the chorionic villi (Figure 4A). Furthermore, expression of the endothelial marker vimentin, which plays an important role in maintaining cell shape and integrity tag, was significantly decreased in SARS-CoV-2-infected samples compared to the uninfected samples (Figure 4B and C, p-value=0.0003). Together, these findings suggest SARS-CoV-2 infection may impact endothelial cell integrity in the placenta even at term.

**Figure 4:**
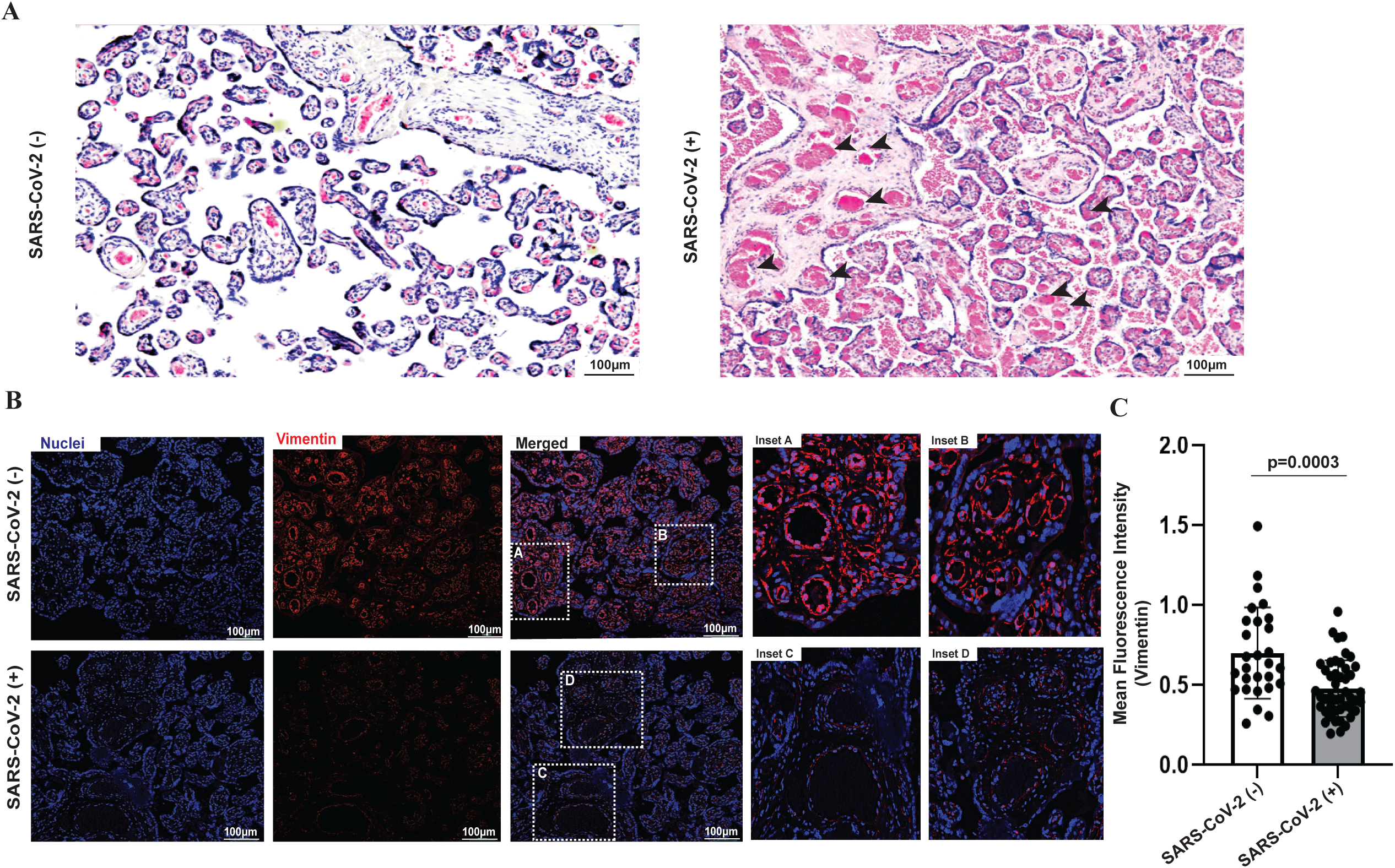
SARS-CoV-2 infection impacts endothelial integrity of the placenta. (A) Foci of chorionic villi with more than ten capillaries in more than ten villi, under objective lens of 10x magnification, characterizing chorangiosis (scale bar represents 100 µm). (B) Expression of vimentin (red) in villous tissue of uninfected (Inset A and B) and infected placentas (Inset C and D) (magnification scale 20X; scale bar represents 100µm). (C) Quantification of mean fluorescence intensity of vimentin in uninfected and SAR-CoV-2 infected placentas, bars represent means and error bars represent standard error of the mean (SEM); *: * p <0.05, ** p < 0.01, ***: p < 0.001; ns: nonsignificant difference.

## 4. Discussion

In this study, our findings reveal several key pathological alterations in the placenta associated with SARS-CoV-2 infection during pregnancy. First, we detected SARS-CoV-2 viral proteins—spike, nucleocapsid, and ORF3a—within the syncytiotrophoblast layer and stroma of the placental villous tissue, indicating active viral replication in the placenta. Additionally, SARS-CoV-2 infection was associated with an increase in syncytial knot formation, a hallmark of placental dysfunction. Notably, placental growth factor (PlGF) expression was significantly reduced in SARS-CoV-2 infected placentas, leading to a higher placental Flt-1/PlGF ratio, primarily driven by lower PlGF levels. This suggests a potential disruption of the local renin-angiotensin system. Furthermore, we observed a decrease in vimentin expression, which is indicative of impaired endothelial integrity. Infected placentas also displayed signs of placental infarctions and excessive fibrin deposits. These findings demonstrate that SARS-CoV-2 infection negatively impacts placental integrity through increased syncytial knots, dysregulated RAS components, and endothelial damage—features commonly associated with preeclampsia. Therefore, these disruptions may underlie the development of a PE-like syndrome in pregnancies complicated by SARS-CoV-2 infection.

Detection of the non-structural accessory protein ORF3a, which is only produced during replication, underscores that SARS-CoV-2 can actively replicate in the placenta. Earlier studies have shown evidence of SARS-CoV-2 virus in STBs, the stromal layer and other compartments of the placenta, supporting our findings [17] Further, a study reported evidence of active *in vivo* viral replication in the placenta after maternal infection in which SARS-CoV-2 replication can increase risk of stillbirths and placentitis [49]. This suggests placental involvement in the pathophysiology of this outcome [49]. Detection of the viral proteins spike, nucleocapsid, and ORF3a suggests that these proteins were persistent through time of delivery when infection occurred much earlier in pregnancy, indicating that the effects of infection could present long after the initial infection has passed. Indeed, persistent viral spike protein in patient plasma and immune cells have been reported suggesting prolonged symptoms post-COVID-19 may stem from constant viral presence and systemic inflammation [50]. Spike protein has been reported in the skull, meninges, and brain samples of acute COVID-19 patients and in post-mortem brain samples [50]. Other SARS-CoV-2 proteins, such as ORF8, have been detected in fetal tissues suggesting transplacental transfer of an accessory protein, potentially contributing to pregnancy complications in both mother and child during infection [51].

Syncytial knots are present in the placenta of normal pregnancies; however, an increase presence is associated with placental maturity and are greatly accelerated in PE [27]. In SARS-CoV-2-infected pregnant women, syncytial knots were significantly higher compared to the uninfected, suggesting premature aging of the placenta. Extensive syncytial knotting is associated with accelerated villous maturation, a histopathological marker for placental sufficiency [29,52]. This accelerated maturation is a manifestation of maternal vascular malperfusion, characterized by small or short hypermature villi for the given gestational period. Studies have shown changes in placental maturity via accelerated maturation and increased risk of maternal vascular malperfusion in SARS-CoV-2-infected placentas [53–55]. Furthermore, syncytial knots have been associated with the transfer of sFlt-1/Flt-1 into the maternal circulation, raising maternal vascular tension and increasing the risk for PE [56]. Our finding that SARS-CoV-2 infection is associated with increased syncytial knot formation and presence of viral proteins in addition to Flt-1 suggest that increased syncytial knots may contribute to the development of a PE-like syndrome in SARS-CoV-2-exposed pregnancies. We also observed placental injury with the presence of placental infarctions, excessive fibrin deposits, and an increase in syncytial nuclear aggregates [12,57– 59], consistent with studies on SARS-CoV-2-infected placental pathology and those seen in the preeclamptic placenta [1,13,36,60]. Further, placental injury caused by SARS-CoV-2 infection is also associated with stillbirth with massive perivillous fibrin deposition, trophoblast necrosis, and chronic histiocytic intervillositis [61], which recapitulate anatomopathological findings in clinical studies and noted in placentas from women with PE [21,57,58,61].

Mounting evidence suggests SARS-CoV-2 can lead to endothelial cell injury and COVID-19 has been recognized as a microvascular and endothelial disease [37,62,63]. SARS-CoV-2 can disrupt the vasculature system in other organs resulting in injury to the kidney, heart, and brain, suggesting that similar disruptions can also occur in the placenta [37,53]. Unvaccinated pregnant women with COVID-19 have been shown to be at increased risk of placental injury due to hypoperfusion and inflammation [16,60]. Studies have shown that SARS-CoV-2 infects endothelial cells in lung tissues via ACE2, increasing endothelial permeability [37,62,64]. Furthermore, higher rates of maternal and fetal vascular malperfusion as a result of SARS-CoV-2 infection are associated with a hypercoagulable state [53]. Indeed, we observed a significant decrease in the expression of vimentin in SARS-CoV-2-infected VT sections compared to uninfected placentas. This suggests potential endothelial injury in the placenta, as further evidenced by the presence of chorangiosis. Such changes may also indicate vascular system disruption, which aligns with our previous findings of decreased PlGF and an elevated Flt-1/PlGF ratio. The vascular dysfunction observed in SARS-CoV-2-infected placentas mirrors endothelial damage seen in PE, where impaired endothelial function is a key feature of the disease. Additionally, SARS-CoV-2 infection has been shown to involve direct interactions between spike proteins and endothelial cells, leading to endothelial dysfunction and microvascular damage in both pulmonary and extrapulmonary systems [65]. Our findings suggest that a similar mechanism may occur in the placenta contributing to a PE-like pathology in SARS-CoV-2-exposed pregnancies.

RAS plays a critical role in regulating blood pressure, hydroelectrolyte balance, and systemic vascular resistance in the kidneys, liver, lungs, and placenta [26,66]. Studies have shown RAS imbalance worsens COVID-19 prognosis, potentially leading to acute respiratory distress syndrome and cytokine storm [13,34]. PlGF plays a key role in normal pregnancy with studies showing that low levels of PlGF mid-pregnancy increase the risks for adverse pregnancy outcomes [67]. In our study, placental Flt-1/PlGF ratio was higher in SARS-CoV-2 infected placenta samples tissue, mainly driven by reduced PlGF levels. We also observed no differences in the levels of AT1-AA in maternal sera between SARS-CoV-2-infected and uninfected pregnant women. This could be due to factors such as antihypertensive medications taken by participants or the timing of sample collection. Interestingly, it has been reported that serum sFlt-1/PlGF levels from infected participants remain unchanged [68], suggesting that SARS-CoV-2 infection may have localized, focal effects in placenta not reflected in serum/plasma biomarkers. Indeed, we observed foci of infection in the placenta, with confocal microscopy revealing localized areas of oxidative stress and DNA damage, colocalized with the SARS-CoV-2 spike protein [69]. These findings indicate that SARS-CoV-2 infection can induce focal placental damage, leaving lasting scars even if infection occurred earlier in pregnancy. This is consistent with a number of recent studies[70]. Furthermore, new mechanistic work from our group suggests that SARS-CoV-2 infection of trophoblast cells in culture or 3D organoids activates secretory autophagy and generates extracellular vesicles carrying ORF3a, which can influence neighboring uninfected areas[71]. This suggests that focal infection can have broader systemic impact. Additionally, persistent SARS-CoV-2 infection may act as a viral reservoir, potentially contributing to long COVID [72]. Growing evidence suggests that pregnant women infected with SARS-CoV-2 may develop long COVID symptoms including cardiovascular conditions[73–75]. Thus, we propose that even limited maternal exposure to SARS-CoV-2, resulting in placental damage, could have broader and longer-term impacts than might be appreciated based on histopathology analysis, which sheds more light than serum markers or symptomology.

Our study is not without limitations. The sample size was insufficient to conduct a secondary analysis comparing pregnant women with hypertensive disorders, both infected and uninfected with SARS-CoV-2. The cohorts included high-risk women with morbidities, such as chronic hypertension, some of whom were taking antihypertensive medications. Additionally, the inability to capture viral strain, infection timing, and severity limits our ability to fully interpret the results. Most samples did not have active infection at the time of delivery or sample collection, and infection could have occurred at any point during pregnancy. Despite these limitations, we observed significant impacts on placental integrity, warranting further studies with larger cohorts and more detailed data on infection timing to clarify the long-term implications of maternal infection on placental health.

In summary, our findings suggest that SARS-CoV-2 infection disrupts placental integrity, with foci of infection in placental villi exacerbating syncytial knot formation, leading to endothelial injury and reduced placental growth factor, potentially contributing to PE-like pathology in infected pregnancies. These results warrant further investigation into the specific locations of viral infection and the long-term implications of maternal infection.

## Acknowledgments and Funding

This work was supported in part by NIH/NICHD grant R01HD091218 to IUM. BRJ was supported by an NIH Institutional National Research Service Award grant [5T32GM136554-03] and by a grant to Baylor College of Medicine from the Howard Hughes Medical Institute through the Gilliam Fellows Program [GT17071]. GMN was supported by Brazilian Coordination of Superior Level Staff Improvement (CAPES) [grant number 88887.712761/2022-00], by CAPES Institutional Internationalization Program (CAPES/PrInt) [grant number 88887.891986/2023-00] and Santander Bank Short-Term International Fellowship [University of Campinas 2023 grant]. MLC was supported by São Paulo Research Foundation (FAPESP) [grant number 2021/09937-1] and by Brazilian National Council for Scientific and Technological Development (CNPq) [grant number 408407/2021-2 and 308378/2022-9]. IUM and MLC were also supported by the Washington University at Saint Louis, USA, McDonnell Academy seed grant for research on infectious diseases and the impact of COVID-19. We thank Robert M. Lawrence for helpful editorial input. We thank all the REBRACO Study Group collaborators, as the core part of the current study.

## Author Contribution Statement

BJ, GMN, MLC and IUM conceptualized the study. BRJ conducted the majority of experiments with help from GMN, DK, and ED. AAT, RTS, JGC, JCK, MLC provided samples. IUM provided conceptual input, edited, and revised the manuscript. All authors revised and approved the final version of the manuscript.

## Informed consent statement

All participants in both cohorts signed an informed consent form authorizing the collection, storage, and use of clinical samples and data.

## Institutional review board statement

Both study cohorts followed all recommended rules for the use of human biological samples, with approval by the respective responsible Research Ethics Committee board. STOP-COVID19 cohort: IRB# 202012075 REBRACO cohort: IRB #31591720.5.0000.5404 issued by the Research Ethics Committee of the University of Campinas, Campinas, São Paulo state, Brazil.

## Data Availability Statement

The data that support the findings of this study are available from the corresponding author upon reasonable request. The data is not publicly available due to privacy or ethical restrictions.

## Declaration of interest

IUM serves on the scientific advisory board of Seed Health. The authors declare that they have no known competing financial interests or personal relationships that could have appeared to influence the work reported in this paper.

### Abbreviations

sFlt-1: soluble fms-like tyrosine kinase-1
Flt1: fms-like tyrosine kinase-1
PlGF: placental growth factor
RAS: renin-angiotensin system
AT1-AA: angiotensin II type-1 receptor autoantibodies
VT: placental villous tissue
PE: Pre-eclampsia
STB: syncytiotrophblast

---

**Supplementary Figure 1:**
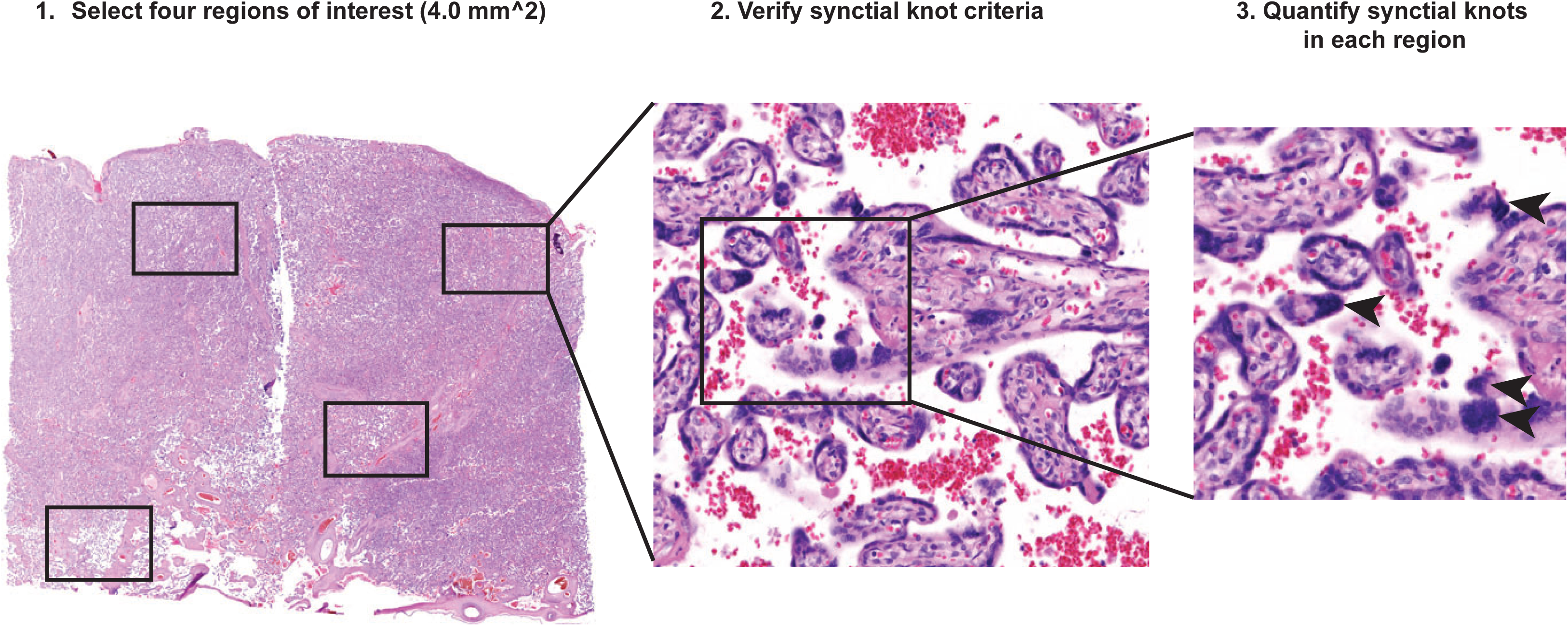
Quantification of syncytial knots (SK). Diagram showing methodology of counting SKs in placental tissue.

**Supplementary Table 1.**
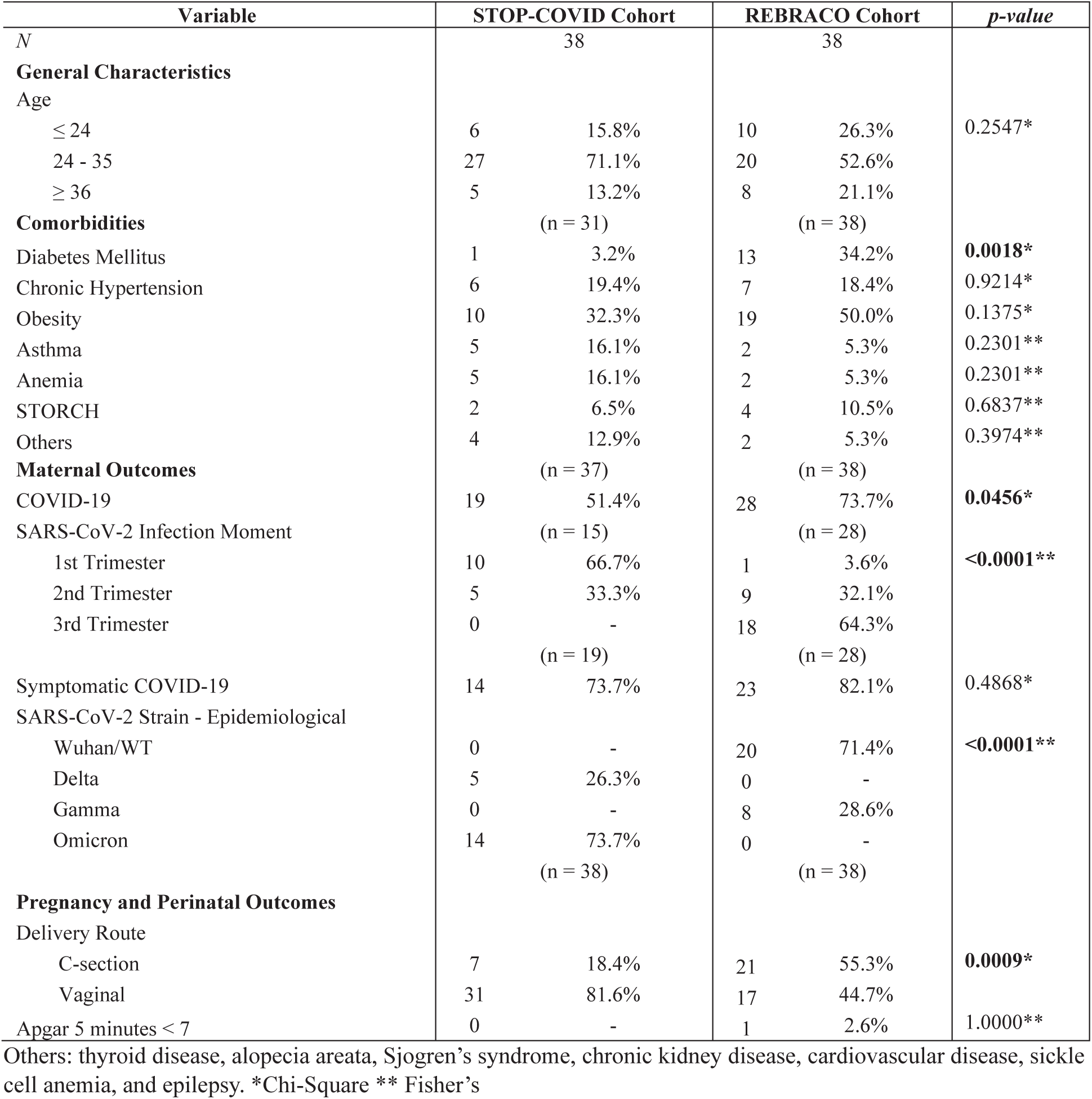
General and clinical characteristics of STOPCOVID-19 and REBRACO cohorts.

**Supplementary Table 2.** Race/ethnicity of STOPCOVID-19 and REBRACO cohorts.

| Variable | STOP-COVID Cohort |  | REBRACO Cohort |  |
| --- | --- | --- | --- | --- |
| N | 38 |  | 38 |  |
| Race | (n = 32) |  | (n= 38) |  |
| White | 17 | 53.1% | 26 | 68.4% |
| Black/African American | 10 | 31.3% | 1 | 2.6% |
| Asian | 4 | 12.5% | 0 | - |
| Latinos | 0 | - | 0 | - |
| Native American | 0 | - | 0 | - |
| Multiracial (*pardo*) | 1 | 3.1% | 11 | 28.9% |

## References

[1] D. Kumar, S. Verma, I.U. Mysorekar, COVID-19 and pregnancy: clinical outcomes; mechanisms, and vaccine efficacy, Translational Research 251 (2023) 84–95. 10.1016/j.trsl.2022.08.007.

[2] R. Sathiya, J. Rajendran, S.S.-M.M. Journal, undefined 2022, COVID-19 and Preeclampsia: Overlapping Features in Pregnancy, Ncbi.Nlm.Nih.Gov (n.d.). https://www.ncbi.nlm.nih.gov/pmc/articles/PMC8798587/ (accessed December 6, 2022).

[3] L.B. Boettcher, T.D. Metz, Maternal and neonatal outcomes following SARS-CoV-2 infection, Semin Fetal Neonatal Med 28 (2023) 101428. 10.1016/j.siny.2023.101428.

[4] C.K.H. Wong, K.T.K. Lau, M.S.H. Chung, I.C.H. Au, K.W. Cheung, E.H.Y. Lau, Y. Daoud, B.J. Cowling, G.M. Leung, Nirmatrelvir/ritonavir use in pregnant women with SARS-CoV-2 Omicron infection: a target trial emulation, Nat Med 30 (2024) 112–116. 10.1038/s41591-023-02674-0.

[5] M. Maher, L. Owens, SARS-CoV-2 infection and female reproductive health: A narrative review, Best Pract Res Clin Endocrinol Metab 37 (2023) 101760. 10.1016/j.beem.2023.101760.

[6] A. Samara, A. Khalil, P. O’Brien, E. Herlenius, The effect of the delta SARS-CoV-2 variant on maternal infection and pregnancy, IScience 25 (2022) 104295. 10.1016/j.isci.2022.104295.

[7] E.L. Reeves, V. Neelam, J.M. Carlson, E.O. Olsen, C.J. Fox, K.R. Woodworth, E. Nestoridi, E. Mobley, S. Montero Castro, P. Dzimira, A. Sokale, L. Sizemore, A.J. Hall, S. Ellington, A. Cohn, S.M. Gilboa, V.T. Tong, Pregnancy and infant outcomes following SARS-CoV-2 infection in pregnancy during delta variant predominance – Surveillance for Emerging Threats to Pregnant People and Infants, Am J Obstet Gynecol MFM 6 (2024) 101265. 10.1016/j.ajogmf.2023.101265.

[8] E.M. Lokken, G.G. Taylor, E.M. Huebner, J. Vanderhoeven, S. Hendrickson, B. Coler, J.S. Sheng, C.L. Walker, S.A. McCartney, N.M. Kretzer, R. Resnick, A. Kachikis, N. Barnhart, V. Schulte, B. Bergam, K.K. Ma, C. Albright, V. Larios, L. Kelley, V. Larios, S. Emhoff, J. Rah, K. Retzlaff, C. Thomas, B.W. Paek, R.J. Hsu, A. Erickson, A. Chang, T. Mitchell, J.K. Hwang, R. Gourley, S. Erickson, S. Delaney, C.R. Kline, K. Archabald, M. Blain, S.M. LaCourse, K.M. Adams Waldorf, Higher severe acute respiratory syndrome coronavirus 2 infection rate in pregnant patients, Am J Obstet Gynecol 225 (2021) 75.e1-75.e16. 10.1016/j.ajog.2021.02.011.

[9] K. Hessami, A.H. Norooznezhad, S. Monteiro, E.R. Barrozo, A.S. Abdolmaleki, S.E. Arian, N. Zargarzadeh, L.S. Shekerdemian, K.M. Aagaard, A.A. Shamshirsaz, COVID-19 Pandemic and Infant Neurodevelopmental Impairment, JAMA Netw Open 5 (2022) e2238941–e2238941. 10.1001/jamanetworkopen.2022.38941.

[10] V. Fajardo-Martinez, F. Ferreira, T. Fuller, M.C. Cambou, T. Kerin, S. Paiola, T. Mok, R. Rao, J. Mohole, R. Paravastu, D. Zhang, P. Marschik, S. Iyer, K. Kesavan, M. da C. Borges Lopes, J.A. A. Britto, M.E. Moreira, P. Brasil, K. Nielsen-Saines, Neurodevelopmental delay in children exposed to maternal SARS-CoV-2 in-utero, Sci Rep 14 (2024) 11851. 10.1038/s41598-024-61918-2.

[11] E.F. Yates, S.B. Mulkey, Viral infections in pregnancy and impact on offspring neurodevelopment: mechanisms and lessons learned, Pediatr Res 96 (2024) 64–72. 10.1038/s41390-024-03145-z.

[12] D.E. Popescu, I. Roșca, A.M.C. Jura, A. Cioca, O. Pop, N. Lungu, Z.-L. Popa, A. Rațiu, M. Boia, Prompt Placental Histopathological and Immunohistochemical Assessment after SARS-CoV-2 Infection during Pregnancy—Our Perspective of a Small Group, Int J Mol Sci 25 (2024) 1836. 10.3390/ijms25031836.

[13] G.M. Nobrega, B.R. Jones, I.U. Mysorekar, M.L. Costa, Preeclampsia in the Context of COVID-19: Mechanisms, Pathophysiology, and Clinical Outcomes, American Journal of Reproductive Immunology 92 (2024). 10.1111/aji.13915.

[14] E.R. Smith, E. Oakley, G.W. Grandner, G. Rukundo, F. Farooq, K. Ferguson, S. Baumann, K.M. Adams Waldorf, Y. Afshar, M. Ahlberg, H. Ahmadzia, V. Akelo, G. Aldrovandi, E. Bevilacqua, N. Bracero, J.S. Brandt, N. Broutet, J. Carrillo, J. Conry, E. Cosmi, F. Crispi, F. Crovetto, M. del Mar Gil, C. Delgado-López, H. Divakar, A.J. Driscoll, G. Favre, I. Fernandez Buhigas, V. Flaherman, C. Gale, C.L. Godwin, S. Gottlieb, E. Gratacós, S. He, O. Hernandez, S. Jones, S. Joshi, E. Kalafat, S. Khagayi, M. Knight, K.L. Kotloff, A. Lanzone, V. Laurita Longo, K. Le Doare, C. Lees, E. Litman, E.M. Lokken, S.A. Madhi, L.A. Magee, R.J. Martinez-Portilla, T.D. Metz, E.S. Miller, D. Money, S. Moungmaithong, E. Mullins, J.B. Nachega, M.C. Nunes, D. Onyango, A. Panchaud, L.C. Poon, D. Raiten, L. Regan, D. Sahota, A. Sakowicz, J. Sanin-Blair, O. Stephansson, M. Temmerman, A. Thorson, S.S. Thwin, B.A. Tippett Barr, J.E. Tolosa, N. Tug, M. Valencia-Prado, S. Visentin, P. von Dadelszen, C. Whitehead, M. Wood, H. Yang, R. Zavala, J.M. Tielsch, Clinical risk factors of adverse outcomes among women with COVID-19 in the pregnancy and postpartum period: a sequential, prospective meta-analysis, Am J Obstet Gynecol 228 (2023) 161–177. 10.1016/j.ajog.2022.08.038.

[15] A. Conde-Agudelo, R. Romero, SARS-CoV-2 infection during pregnancy and risk of preeclampsia: a systematic review and meta-analysis, Am J Obstet Gynecol 226 (2022) 68–89.e3. 10.1016/j.ajog.2021.07.009.

[16] S. Verma, E.B. Carter, I.U. Mysorekar, SARS-CoV2 and pregnancy: An invisible enemy?, American Journal of Reproductive Immunology 84 (2020). 10.1111/aji.13308.

[17] S. Verma, C.S. Joshi, R.B. Silverstein, M. He, E.B. Carter, I.U. Mysorekar, SARS-CoV-2 colonization of maternal and fetal cells of the human placenta promotes alteration of local renin-angiotensin system, Med 2 (2021) 575–590.e5. 10.1016/j.medj.2021.04.009.

[18] N.N. Mahajan, S. Kesarwani, P. Kumbhar, P. Kuppusamy, M. Pophalkar, P. Thamke, R. Asawa, S. Sharan, S.D. Mahale, R.K. Gajbhiye, Increased risk of early-onset preeclampsia in pregnant women with COVID-19, Hypertens Pregnancy 42 (2023). 10.1080/10641955.2023.2187630.

[19] R. Gholami, N. Borumandnia, E. Kalhori, M. Taheri, N. Khodakarami, The impact of covid-19 pandemic on pregnancy outcome, BMC Pregnancy Childbirth 23 (2023) 811. 10.1186/s12884-023-06098-z.

[20] E.A. Phipps, R. Thadhani, T. Benzing, S.A. Karumanchi, Pre-eclampsia: pathogenesis, novel diagnostics and therapies, Nat Rev Nephrol 15 (2019) 275–289. 10.1038/s41581-019-0119-6.

[21] M. Mendoza, I. Garcia-Ruiz, N. Maiz, C. Rodo, P. Garcia-Manau, B. Serrano, R.M. Lopez-Martinez, J. Balcells, N. Fernandez-Hidalgo, E. Carreras, A. Suy, Pre-eclampsia-like syndrome induced by severe COVID-19: a prospective observational study, BJOG 127 (2020) 1374–1380. 10.1111/1471-0528.16339.

[22] S.Q. Wei, M. Bilodeau-Bertrand, S. Liu, N. Auger, The impact of COVID-19 on pregnancy outcomes: a systematic review and meta-analysis, Can Med Assoc J 193 (2021) E540–E548. 10.1503/cmaj.202604.

[23] M. Abedzadeh-Kalahroudi, M. Sehat, Z. Vahedpour, P. Talebian, Maternal and neonatal outcomes of pregnant patients with COVID-19: A prospective cohort study, International Journal of Gynecology & Obstetrics 153 (2021) 449–456. 10.1002/ijgo.13661.

[24] M.W. Stark, L. Clark, R.D. Craver, Histologic Differences in Placentas of Preeclamptic/Eclamptic Gestations by Birthweight, Placental Weight, and Time of Onset, Pediatric and Developmental Pathology 17 (2014) 181–189. 10.2350/13-09-1378-OA.1.

[25] C.S. Roland, J. Hu, C.-E. Ren, H. Chen, J. Li, M.S. Varvoutis, L.W. Leaphart, D.B. Byck, X. Zhu, S.-W. Jiang, Morphological changes of placental syncytium and their implications for the pathogenesis of preeclampsia, Cellular and Molecular Life Sciences 73 (2016) 365–376. 10.1007/s00018-015-2069-x.

[26] L. Yart, E. Roset Bahmanyar, M. Cohen, B. Martinez de Tejada, Role of the Uteroplacental Renin–Angiotensin System in Placental Development and Function, and Its Implication in the Preeclampsia Pathogenesis, Biomedicines 9 (2021) 1332. 10.3390/biomedicines9101332.

[27] A. Rajakumar, A.S. Cerdeira, S. Rana, Z. Zsengeller, L. Edmunds, A. Jeyabalan, C.A. Hubel, I.E. Stillman, S.M. Parikh, S.A. Karumanchi, Transcriptionally Active Syncytial Aggregates in the Maternal Circulation May Contribute to Circulating Soluble Fms-Like Tyrosine Kinase 1 in Preeclampsia, Hypertension 59 (2012) 256–264. 10.1161/HYPERTENSIONAHA.111.182170.

[28] Y.P. Wong, F.C. Cheah, K.K. Wong, S.A. Shah, S.E. Phon, B.K. Ng, P.S. Lim, T.Y. Khong, G.C. Tan, Gardnerella vaginalis infection in pregnancy: Effects on placental development and neonatal outcomes, Placenta 120 (2022) 79–87. 10.1016/j.placenta.2022.02.018.

[29] V. Taché, D.Y. LaCoursiere, A. Saleemuddin, M.M. Parast, Placental expression of vascular endothelial growth factor receptor-1/soluble vascular endothelial growth factor receptor-1 correlates with severity of clinical preeclampsia and villous hypermaturity, Hum Pathol 42 (2011) 1283–1288. 10.1016/j.humpath.2010.11.018.

[30] K. Loukeris, R. Sela, R.N. Baergen, Syncytial Knots as a Reflection of Placental Maturity: Reference Values for 20 to 40 Weeks’ Gestational Age, Pediatric and Developmental Pathology 13 (2010) 305–309. 10.2350/09-08-0692-OA.1.

[31] S. Bernardi, F. Tonon, M. Barbieri, G. Zamagni, R. Nuredini, L. Perer, S. Comar, B. Toffoli, L. Ronfani, G. Ricci, B. Fabris, T. Stampalija, A longitudinal study on the effect of obesity upon circulating renin-angiotensin system in normal pregnancy, Nutrition, Metabolism and Cardiovascular Diseases 34 (2024) 771–782. 10.1016/j.numecd.2023.10.030.

[32] G.M. Nobrega, J.P. Guida, J.M. Novaes, L.M. Solda, L. Pietro, A.G. Luz, G.J. Lajos, C.C. Ribeiro-do-Valle, R.T. Souza, J.G. Cecatti, I.U. Mysorekar, T.Z. Dias, M. Laura Costa, Role of biomarkers (sFlt-1/PlGF) in cases of COVID-19 for distinguishing preeclampsia and guiding clinical management, Pregnancy Hypertens 31 (2023) 32–37. 10.1016/j.preghy.2022.11.008.

[33] P.E. Andronikidi, E. Orovou, E. Mavrigiannaki, V. Athanasiadou, M. Tzitiridou-Chatzopoulou, G. Iatrakis, E. Grapsa, Placental and Renal Pathways Underlying Pre-Eclampsia, Int J Mol Sci 25 (2024) 2741. 10.3390/ijms25052741.

[34] S.J. Delforce, E.R. Lumbers, S.J. Ellery, P. Murthi, K.G. Pringle, Dysregulation of the placental renin–angiotensin system in human fetal growth restriction, Reproduction 158 (2019) 237–245. 10.1530/REP-18-0633.

[35] C.A. Penfield, S.G. Brubaker, M.A. Limaye, J. Lighter, A.J. Ratner, K.M. Thomas, J.A. Meyer, A.S. Roman, Detection of severe acute respiratory syndrome coronavirus 2 in placental and fetal membrane samples, Am J Obstet Gynecol MFM 2 (2020) 100133. 10.1016/j.ajogmf.2020.100133.

[36] A. Antolini-Tavares, G.M. Nobrega, J.P. Guida, A.G. Luz, G.J. Lajos, Carolina C.R. do-Valle, R.T. Souza, J.G. Cecatti, I.U. Mysorekar, M.L. Costa, Morphological placental findings in women infected with SARS-CoV-2 according to trimester of pregnancy and severity of disease, Placenta 139 (2023) 190–199. 10.1016/j.placenta.2023.06.015.

[37] S. Xu, I. Ilyas, J. Weng, Endothelial dysfunction in COVID-19: an overview of evidence, biomarkers, mechanisms and potential therapies, Acta Pharmacol Sin 44 (2023) 695–709. 10.1038/s41401-022-00998-0.

[38] J. Meng, G. Xiao, J. Zhang, X. He, M. Ou, J. Bi, R. Yang, W. Di, Z. Wang, Z. Li, H. Gao, L. Liu, G. Zhang, Renin-angiotensin system inhibitors improve the clinical outcomes of COVID-19 patients with hypertension, Emerg Microbes Infect 9 (2020) 757–760. 10.1080/22221751.2020.1746200.

[39] M.J. Lim, S. Lakshminrusimha, H. Hedriana, T. Albertson, Pregnancy and Severe ARDS with COVID-19: Epidemiology, Diagnosis, Outcomes and Treatment, Semin Fetal Neonatal Med 28 (2023) 101426. 10.1016/j.siny.2023.101426.

[40] A.I. Rodriguez-Perez, C.M. Labandeira, M.A. Pedrosa, R. Valenzuela, J.A. Suarez-Quintanilla, M. Cortes-Ayaso, P. Mayán-Conesa, J.L. Labandeira-Garcia, Autoantibodies against ACE2 and angiotensin type-1 receptors increase severity of COVID-19, J Autoimmun 122 (2021) 102683. 10.1016/j.jaut.2021.102683.

[41] K. Kuba, T. Yamaguchi, J.M. Penninger, Angiotensin-Converting Enzyme 2 (ACE2) in the Pathogenesis of ARDS in COVID-19, Front Immunol 12 (2021). 10.3389/fimmu.2021.732690.

[42] S. Rysz, J. Al-Saadi, A. Sjöström, M. Farm, F. Campoccia Jalde, M. Plattén, H. Eriksson, M. Klein, R. Vargas-Paris, S. Nyrén, G. Abdula, R. Ouellette, T. Granberg, M. Jonsson Fagerlund, J. Lundberg, COVID-19 pathophysiology may be driven by an imbalance in the renin-angiotensin-aldosterone system, Nat Commun 12 (2021) 2417. 10.1038/s41467-021-22713-z.

[43] K. Chau, A. Hennessy, A. Makris, Placental growth factor and pre-eclampsia, J Hum Hypertens 31 (2017) 782–786. 10.1038/jhh.2017.61.

[44] C.R. Palma Dos Reis, J. O’Sullivan, E.O. Ohuma, T. James, A.T. Papageorghiou, M. Vatish, A.S. Cerdeira, The ratio of soluble fms-like tyrosine kinase 1 to placental growth factor predicts time to delivery and mode of birth in patients with suspected preeclampsia: a secondary analysis of the INSPIRE trial, Am J Obstet Gynecol 232 (2025) 317.e1-317.e17. 10.1016/j.ajog.2024.06.010.

[45] M.C. Sharps, D.J.L. Hayes, S. Lee, Z. Zou, C.A. Brady, Y. Almoghrabi, A. Kerby, K.K. Tamber, C.J. Jones, K.M. Adams Waldorf, A.E.P. Heazell, A structured review of placental morphology and histopathological lesions associated with SARS-CoV-2 infection, Placenta 101 (2020) 13–29. 10.1016/j.placenta.2020.08.018.

[46] R.T. Souza, J.G. Cecatti, R.C. Pacagnella, C.C. Ribeiro-Do-Valle, A.G. Luz, G.J. Lajos, G.M. Nobrega, T.B. Griggio, C.M. Charles, S.F. Bento, C. Silveira, F.G. Surita, M.J. Miele, R.P. Tedesco, K.G. Fernandes, S.H.A. Martins-Costa, F.J.A. Peret, F.E. Feitosa, R. Mattar, E. Traina, E. V. Cunha Filho, J. Vettorazzi, S.M. Haddad, C.B. Andreucci, J.P. Guida, M.D. Correa Junior, M.A.B. Dias, L. De Oliveira, E.F. Melo Junior, M.G.Q. Luz, M.L. Costa, R.T. Souza, M.L. Costa, C.C. Ribeiro-do-Valle, A.G. Luz, G.J. Lajos, G.M. Nobrega, T.B. Griggrio, C.M. Charles, S.F. Bento, C. Silveira, F.G. Surita, M.J. Miele, S. Metelus, L. Castro, S. Pabon, A.D. Silva, P.S.R. Junior, T.G. Sardinha, R.R. Japenga, E.R.F. Urquiza, M.R. Machado, M.M. Simões, L.M. Solda, J.V. Freitas-Jesus, R.E. Soeiro, R.P. Tedesco, K.G. Fernandes, P.B. Peres, C.L. Arbeli, R.M. Quevedo, C.F. Yamashita, J.D. Corradin, I. Bergamini, S.H.A. Martins-Costa, J.G.L. Ramos, M.L.R. Oppermann, L.S. Quadro, L. Marins, É. V. Paniz, T.V. Xavier, F.J.A. Peret, M.H.L. Almeida, B.F.V. Moura, L.R. França, H. Vieira, R.B. Aquino, A.C. Costa, F.E. Feitosa, D. Pinheiro, D. Cordeiro, P.L. Miná, C. Dornellas, R. Mattar, E. Traina, S. Yazaki-Sun, P. Mota, A.C. Soares, E. V. Cunha Filho, J. Vettorazzi, E. Machado, A. Bergmann, G.R. Santos, S.M. Haddad, A. Tosetto, S. Savazoni, C.B. Andreucci, B.E. Parreira, J.P. Guida, M.D. Correa Junior, C. Leal, R. Amana, M.A.B. Dias, M. Nakamura-Pereira, B.O. Guerra, G. Gorga, L. De Oliveira, K.F.A. Oliveira, M.E.V. Makyama, E.F. Melo Junior, D.F. Leite, I. Monteiro, M.G.Q. Luz, I.R. Pereira, C.A. Salustrino, V.B. Pontes, R.A. Silva Franco, J.P. Bilibio, G.P.F. Brito, H.P.C. Pinto, D.L. Oliveira, A.A. Guerra, A.O. Moura, N. Pantoja, F. David, A. Silva, The COVID-19 pandemic in Brazilian pregnant and postpartum women: results from the REBRACO prospective cohort study, Sci Rep 12 (2022). 10.1038/S41598-022-15647-Z,.

[47] A. Dantas-Silva, F. Garanhani Surita, R. Souza, L. Rocha, J. Paulo Guida, R. Pacagnella, R. Tedesco, K. Fernandes, S. Martins-Costa, F. Peret, F. Feitosa, E. Traina, E. Cunha Filho, J. Vettorazzi, S. Haddad, C. Andreucci, M. Correa Junior, M. Dias, L. de Oliveira, E. Melo Junior, M. Luz, J. Guilherme Cecatti, M. Laura Costa, A. Fleming, C.C. Ribeiro Do-Valle, A.G. Luz, G.J. Lajos, G.M. Nobrega, T.B. Griggio, C.M. Charles, S.F. Bento, C. Silveira, M.J. Miele, L. Bahamondes, S. Metelus, L. Castro, S. Pabon, R. Esteves Soeiro, A. Antolini, P.S. R Junior, T.G. Sardinha, R.R. Japenga, E.R. F Urquiza, M.R. Machado, M. Maria Simões, L.M. Solda, S. Yazaki-Sun, P. Mota, A.C. Soares, E. Machado, A. Bergmann, G. Raupp dos Santos, P.B. Peres, C.L. Arbeli, R.M. Quevedo, C.F. Yamashita, J.D. Corradin, I. Bergamini, J.L. Geraldo Ramos, M.R. Lúcia Oppermann, L.S. Quadro, L. Marins, É. V Paniz, T. Vicentini Xavier, B.E. Parreira, A.M. Tosetto, S.O. Savazoni, A.C. Costa, M.H. Almeida, B.F. Moura, L.R. França, H. Vieira, R.B. Aquino, D.F. Leite, I. Monteiro, M. Nakamura-Pereira, B.O. Guerra, G. Gorga, D. Pinheiro, D. Cordeiro, P.L. Miná, C. Dornellas, K.F. Oliveira, M. Emi Varicoda Makyama, C. Leal, R. Amana, C.O. Santos, M.M. dos Santos, C. Neto, T. Gomes, I.R. Pereira, C. Andrade Salustrino, V.B. Pontes, R. Allen da Silva Franco, J. Paolo Bilibio, G.P. F Brito, H.C. Paula Pinto, D. Leal de Oliveira, A.A. Guerra, A.O. Moura, N. Pantoja, F. David, A. Silva, J. Vasconcellos Freitas-Jesus, A.M. Bacha, A. Borovac-Pinheiro, B.G. Pereira, E.M. Amaral, E. Ferreira, H. MBPM Milanez, J.P. S Caldas, L.F. Baccaro, M. Nomura, P.M. Rehder, R.Z. Simone, R. Passini Jr, C. Torrezan, J.L. P Modena, M.N. Nunes dos Santos, S.T. M Marba, T.R. Zumpano dos Santos, Brazilian black women are at higher risk for COVID-19 complications: an analysis of REBRACO, a national cohort, SciELO BrasilA Dantas-Silva, FG Surita, R Souza, L Rocha, JP Guida, R Pacagnella, R TedescoRevista Brasileira de Ginecologia e Obstetrícia, 2023•SciELO Brasil 45 (2023) 253–260. 10.1055/s-0043-1770133.

[48] G.M. Nobrega, E.R. McColl, A. Antolini-Tavares, R.T. Souza, J.G. Cecatti, M.L. Costa, I.U. Mysorekar, Placentas From SARS-CoV-2 Infection During Pregnancy Exhibit Foci of Oxidative Stress and DNA Damage, American Journal of Reproductive Immunology 93 (2025). 10.1111/aji.70034.

[49] A.-P. Radan, P. Renz, L. Raio, A.-S. Villiger, V. Haesler, M. Trippel, D. Surbek, SARS-CoV-2 replicates in the placenta after maternal infection during pregnancy, Front Med (Lausanne) 11 (2024). 10.3389/fmed.2024.1439181.

[50] Z. Rong, H. Mai, G. Ebert, S. Kapoor, V.G. Puelles, J. Czogalla, S. Hu, J. Su, D. Prtvar, I. Singh, J. Schädler, C. Delbridge, H. Steinke, H. Frenzel, K. Schmidt, C. Braun, G. Bruch, V. Ruf, M. Ali, K.-W. Sühs, M. Nemati, F. Hopfner, S. Ulukaya, D. Jeridi, D. Mistretta, Ö.S. Caliskan, J.M. Wettengel, F. Cherif, Z.I. Kolabas, M. Molbay, I. Horvath, S. Zhao, N. Krahmer, A.Ö. Yildirim, S. Ussar, J. Herms, T.B. Huber, S. Tahirovic, S.M. Schwarzmaier, N. Plesnila, G. Höglinger, B. Ondruschka, I. Bechmann, U. Protzer, M. Elsner, H.S. Bhatia, F. Hellal, A. Ertürk, Persistence of spike protein at the skull-meninges-brain axis may contribute to the neurological sequelae of COVID-19, Cell Host Microbe 32 (2024) 2112–2130.e10. 10.1016/j.chom.2024.11.007.

[51] T. Azamor, D. Familiar-Macedo, G.M. Salem, C. Onwubueke, I. Melano, L. Bian, Z. Vasconcelos, K. Nielsen-Saines, X. Wu, J.U. Jung, F. Lin, O. Goje, E. Chien, S. Gordon, C.B. Foster, H. Aly, R.M. Farrell, W. Chen, S.-S. Foo, Transplacental SARS-CoV-2 protein ORF8 binds to complement C1q to trigger fetal inflammation, EMBO J 43 (2024) 5494–5529. 10.1038/s44318-024-00260-9.

[52] Y. Ma, L. Kong, Q. Ge, Y. Lu, M. Hong, Y. Zhang, C. Ruan, P. Gao, Complement 5a-mediated trophoblasts dysfunction is involved in the development of pre-eclampsia, J Cell Mol Med 22 (2018) 1034–1046. 10.1111/jcmm.13466.

[53] T. Le, D. Lee, L.S. Brown, B.W. Payton, P. Sepulveda, J. Sisman, R.L. Leon, L.F. Chalak, I.N. Mir, Placental pathology in SARS-CoV-2 infected pregnancies: A single-institution retrospective cohort analysis, J Neonatal Perinatal Med 17 (2024) 623–636. 10.3233/NPM-230177.

[54] D. Kidron, I. Vainer, Y. Fisher, R. Sharony, Automated image analysis of placental villi and syncytial knots in histological sections, Placenta 53 (2017) 113–118. 10.1016/j.placenta.2017.04.004.

[55] T.Y. Khong, E.E. Mooney, I. Ariel, N.C.M. Balmus, T.K. Boyd, M.-A. Brundler, H. Derricott, M.J. Evans, O.M. Faye-Petersen, J.E. Gillan, A.E.P. Heazell, D.S. Heller, S.M. Jacques, S. Keating, P. Kelehan, A. Maes, E.M. McKay, T.K. Morgan, P.G.J. Nikkels, W.T. Parks, R.W. Redline, I. Scheimberg, M.H. Schoots, N.J. Sebire, A. Timmer, G. Turowski, J.P. van der Voorn, I. van Lijnschoten, S.J. Gordijn, Sampling and Definitions of Placental Lesions: Amsterdam Placental Workshop Group Consensus Statement, Arch Pathol Lab Med 140 (2016) 698–713. 10.5858/arpa.2015-0225-CC.

[56] H. Zeisler, E. Llurba, F. Chantraine, M. Vatish, A.C. Staff, M. Sennström, M. Olovsson, S.P. Brennecke, H. Stepan, D. Allegranza, P. Dilba, M. Schoedl, M. Hund, S. Verlohren, Predictive Value of the sFlt-1:PlGF Ratio in Women with Suspected Preeclampsia, New England Journal of Medicine 374 (2016) 13–22. 10.1056/NEJMoa1414838.

[57] A.C.M. Nascimento, E. Avvad-Portari, M. Meuser-Batista, T.C. Conde, R.A.M. de Sá, N. Salomao, K. Rabelo, E.S. Ciasca, M. de Oliveira Brendolin, Z. Vasconcelos, P. Brasil, M.E. Moreira, Histopathological and clinical analysis of COVID-19-infected placentas, Surgical and Experimental Pathology 7 (2024) 4. 10.1186/s42047-024-00146-4.

[58] M.A. Snow, M.K. Annavajhala, S.Z. Moscovitz, A.-C. Uhlemann, L. Debelenko, Placental Infection with Different SARS-CoV-2 Variants Leading to Stillbirth: Report of Two Cases, COVID 5 (2025) 8. 10.3390/covid5010008.

[59] J. Li, Y. Zhou, J. Ma, Q. Zhang, J. Shao, S. Liang, Y. Yu, W. Li, C. Wang, The long-term health outcomes, pathophysiological mechanisms and multidisciplinary management of long COVID, Signal Transduct Target Ther 8 (2023) 416. 10.1038/s41392-023-01640-z.

[60] R. Di Girolamo, A. Khalil, S. Alameddine, E. D’Angelo, C. Galliani, B. Matarrelli, D. Buca, M. Liberati, G. Rizzo, F. D’Antonio, Placental histopathology after SARS-CoV-2 infection in pregnancy: a systematic review and meta-analysis, Am J Obstet Gynecol MFM 3 (2021) 100468. 10.1016/j.ajogmf.2021.100468.

[61] A. Li, D.A. Schwartz, A. Vo, R. VanAbel, C. Coler, E. Li, B. Lukman, B. Del Rosario, A. Vong, M. Li, K.M. Adams Waldorf, Impact of SARS-CoV-2 infection during pregnancy on the placenta and fetus, Semin Perinatol 48 (2024) 151919. 10.1016/j.semperi.2024.151919.

[62] R. Kumar, Ö. Aktay-Cetin, V. Craddock, D. Morales-Cano, D. Kosanovic, A. Cogolludo, F. Perez-Vizcaino, S. Avdeev, A. Kumar, A.K. Ram, S. Agarwal, A. Chakraborty, R. Savai, V. de Jesus Perez, B.B. Graham, G. Butrous, N.K. Dhillon, Potential long-term effects of SARS-CoV-2 infection on the pulmonary vasculature: Multilayered cross-talks in the setting of coinfections and comorbidities, PLoS Pathog 19 (2023) e1011063–e1011063. 10.1371/journal.ppat.1011063.

[63] X. Wu, M. Xiang, H. Jing, C. Wang, V.A. Novakovic, J. Shi, Damage to endothelial barriers and its contribution to long COVID, Angiogenesis 27 (2024) 5–22. 10.1007/s10456-023-09878-5.

[64] I. Bernard, D. Limonta, L. Mahal, T. Hobman, Endothelium Infection and Dysregulation by SARS-CoV-2: Evidence and Caveats in COVID-19, Viruses 13 (2020) 29. 10.3390/v13010029.

[65] L. Perico, A. Benigni, G. Remuzzi, SARS-CoV-2 and the spike protein in endotheliopathy, Trends Microbiol 32 (2024) 53–67. 10.1016/j.tim.2023.06.004.

[66] C.R.V. Leal, L.B. Costa, G.C. Ferreira, A. de M. Ferreira, F.M. Reis, A.C. Simões e Silva, Renin-angiotensin system in normal pregnancy and in preeclampsia: A comprehensive review, Pregnancy Hypertens 28 (2022) 15–20. 10.1016/j.preghy.2022.01.011.

[67] R.A. Gladstone, S. Ahmed, E. Huszti, K. McLaughlin, J.W. Snelgrove, J. Taher, S.R. Hobson, R.C. Windrim, K.E. Murphy, J.C. Kingdom, Midpregnancy Placental Growth Factor Screening and Early Preterm Birth, JAMA Netw Open 7 (2024) e2444454–e2444454. 10.1001/jamanetworkopen.2024.44454.

[68] G.M. Nobrega, L. Pietro, S.L. Dariva, I.A. Vasconcelos-Silva, M.P. Manari, B. Polli, A.B. Simões, J.S. de Almeida, R. Moschetta, C.C. Ribeiro-do-Valle, J.P. Siqueira Guida, R.T. Souza, J.G. Cecatti, I.U. Mysorekar, A.S. Picoloto, M.L. Costa, Preeclampsia biomarkers (sFlt-1/PlGF) dynamics are not disrupted by SARS-CoV-2 infection during pregnancy in a hypertensive disorder SARS-CoV-2 vaccinated cohort, Pregnancy Hypertens 39 (2025) 101196. 10.1016/j.preghy.2025.101196.

[69] G.M. Nobrega, B.R. Jones, I.U. Mysorekar, M.L. Costa, Preeclampsia in the Context of COVID-19: Mechanisms, Pathophysiology, and Clinical Outcomes, American Journal of Reproductive Immunology 92 (2024). 10.1111/AJI.13915.

[70] L.C. Gabby, C.K. Jones, B.B. McIntyre, Z. Manalo, M. Meads, D.P. Pizzo, J. Diaz-Vigil, F. Soncin, K.M. Fisch, G.A. Ramos, M.B. Jacobs, M.M. Parast, Chronic villitis as a distinctive feature of placental injury in maternal SARS-CoV-2 infection, Am J Obstet Gynecol 232 (2024). 10.1016/j.ajog.2024.04.002.

[71] D. Kumar, R.M. Karvas, B. Jones, E. McColl, E. Diveley, S. Jash, S. Sharma, J. Kelly, T. Theunissen, I.U. Mysorekar, SARS-CoV-2 ORF3a Protein Impairs Syncytiotrophoblast Maturation, Alters ZO-1 Localization, and Shifts Autophagic Pathways in Trophoblast Cells and 3D Organoids, BioRxiv (2024). 10.1101/2024.09.25.614931.

[72] M. Ghafari, M. Hall, T. Golubchik, D. Ayoubkhani, T. House, G. MacIntyre-Cockett, H.R. Fryer, L. Thomson, A. Nurtay, S.A. Kemp, L. Ferretti, D. Buck, A. Green, A. Trebes, P. Piazza, L.J. Lonie, R. Studley, E. Rourke, D.L. Smith, M. Bashton, A. Nelson, M. Crown, C. McCann, G.R. Young, R.A.N. dos Santos, Z. Richards, M.A. Tariq, R. Cahuantzi, J. Barrett, C. Fraser, D. Bonsall, A.S. Walker, K. Lythgoe, Prevalence of persistent SARS-CoV-2 in a large community surveillance study, Nature 2024 626:8001 626 (2024) 1094–1101. 10.1038/s41586-024-07029-4.

[73] H. Kandemir, G.A. Bülbül, E. Kirtiş, S. Güney, C.Y. Sanhal, İ.İ. Mendilcioğlu, Evaluation of long-COVID symptoms in women infected with SARS-CoV-2 during pregnancy, International Journal of Gynecology and Obstetrics 164 (2024) 148–156. 10.1002/IJGO.14972.

[74] C. Zang, D. Guth, A.M. Bruno, Z. Xu, H. Li, N. Ammar, R. Chew, N. Guthe, E. Hadley, R. Kaushal, T. Love, B.M. McGrath, R.C. Patel, E.C. Seibert, Y. Senathirajah, S.K. Singh, F. Wang, M.G. Weiner, K.J. Wilkins, Y. Zhang, T.D. Metz, E. Hill, T.W. Carton, Long COVID after SARS-CoV-2 during pregnancy in the United States, Nature Communications 2025 16:1 16 (2025) 1–14. 10.1038/s41467-025-57849-9.

[75] S. Su, Y. Zhao, N. Zeng, X. Liu, Y. Zheng, J. Sun, Y. Zhong, S. Wu, S. Ni, Y. Gong, Z. Zhang, N. Gao, K. Yuan, W. Yan, L. Shi, A. V. Ravindran, T. Kosten, J. Shi, Y. Bao, L. Lu, Epidemiology, clinical presentation, pathophysiology, and management of long COVID: an update, Molecular Psychiatry 2023 28:10 28 (2023) 4056–4069. 10.1038/s41380-023-02171-3.

